# The phytonutrient cinnamaldehyde limits intestinal inflammation and enteric parasite infection

**DOI:** 10.1101/2021.03.02.433624

**Authors:** Ling Zhu, Audrey I.S. Andersen-Civil, Laura J. Myhill, Stig M. Thamsborg, Witold Kot, Lukasz Krych, Dennis S. Nielsen, Alexandra Blanchard, Andrew R. Williams

## Abstract

Phytonutrients such as cinnamaldehyde (CA) have been studied for their effects on metabolic diseases, but their influence on mucosal inflammation and immunity to enteric infection are not well documented. Here, we show that consumption of CA significantly down-regulates transcriptional pathways connected to inflammation in the small intestine of mice. During infection with the enteric helminth Heligomosomoides polygyrus, CA-treated mice displayed higher growth rates and less worms, concomitant with altered T-cell populations in mesenteric lymph nodes. Furthermore, infection-induced changes in gene pathways connected to cell cycle and mitotic activity were counteracted by CA. Mechanically, CA did not appear to exert activity through a prebiotic effect, as CA treatment did not significantly change the composition of the gut microbiota. Instead, in vitro experiments showed that CA directly induced xenobiotic metabolizing pathways in intestinal epithelial cells and suppressed endotoxin-induced inflammatory responses in macrophages. Thus, CA down-regulates inflammatory pathways in the intestinal mucosa and regulates host responses to enteric infection. These properties appear to be largely independent of the gut microbiota and instead connected to CA’s ability to induce antioxidant pathways in intestinal cells. Our results encourage further investigation into the use of CA and related phytonutrients as functional food components to promote intestinal health in humans and animals.

## 1. Introduction

Dietary components can play an important role in the regulation of immune function and inflammation in the gastrointestinal tract, and the severity of chronic inflammatory diseases, such as colitis and diabetes, may be modulated by diet [1]. Dietary compounds continuously pass through the gut, leading to close contact with the intestinal epithelium and the commensal gut microbiota (GM). In this way, they may induce either direct immunomodulatory effects on mucosal immune cells, or indirectly modulate immune responses due to prebiotic modification of the GM and production of microbial-derived metabolites. For example, some dietary contents may be fermented by commensal bacteria, resulting in the production metabolites such as short chain fatty acids, which can affect metabolic and immune function such as the production of regulatory T cells and luminal IgA [2, 3].

Phytonutrients are defined as plant compounds that, when consumed as part of the diet, have beneficial effects on health and/or disease. A number of studies have reported that dietary phytonutrients can regulate mucosal barrier function and intestinal microbiota to regulate inflammatory processes in human and animal health [4]. For example, proanthocyanidins (common plant secondary metabolites found in berries) attenuate intestinal permeability and oxidative stress induced by lipopolysaccharide (LPS) in rats [5–7]. Cinnamaldehyde (CA) is a bioactive compound that is the major component of the essential oil from cinnamon, and responsible for its distinctive aroma. It is thus commonly used in the food industry as a flavouring agent and as a natural preservative [8]. In addition, it has attracted increasing interest for its anti-inflammatory properties. CA has been reported to suppress the LPS-induced activation of NF-κB and oligomerization of toll-like receptor 4 (TLR4) in mouse macrophages *in vitro*, showing the potential mechanism for anti-inflammatory activity [9]. Consistent with this, CA has shown promise in alleviating chemically-induced colitis in mice, as well as playing a beneficial role in the regulation of obesity and chronic, low-grade inflammation [10–12].

Despite these putative beneficial effects of CA and other phytonutrients on metabolic diseases, effects on mucosal immunity and responses to infectious pathogens have been studied in less detail. CA is increasingly used as a feed additive in livestock production, where it may increase levels of serum immunoglobulin and circulating lymphocytes [13, 14]. Moreover, chickens fed CA are more resistant to infection with the protozoan parasite *Eimeria tenella*, and have higher levels of specific antibodies following infection [15]. Thus, CA and related phytonutrients may play a role in promoting gut health and increasing resistance to enteric pathogens, which is of critical importance for sustainable animal and human health in an era of widespread antimicrobial drug resistance.

Parasitic worms (helminths) are one of the most widespread infectious agents worldwide. In humans, helminths infect more than a billion people, causing substantial morbidity, and infection in livestock reduces animal performance and results in significant economic penalties for farmers [16]. Moreover, helminth infections induce multi-faceted mucosal immune responses, incorporating both pro-inflammatory as well as type-2 and modified regulatory responses that resemble intestinal diseases such as food allergies and ulcerative colitis [17]. Nutritional interventions that can reduce pathology, and perhaps also favour the development of protective immune mechanisms, would be of great benefit for sustainable control of parasitic infection. We have previously reported that dietary CA enhances acquisition of parasite-specific antibodies in pigs following infection with the helminth *Ascaris suum*, whilst suppressing the expression of genes encoding inflammatory cytokines [18]. Our objective here was to explore in more detail how CA exerts immunoregulatory effects in resistance to infectious disease. Experimental mouse models have been proven highly reliable and indispensable to understand the mechanisms of intestinal disorders [19]. We utilized the enteric helminth *Heligomosomoides polygyrus*, a well-characterized murine model of human and livestock helminths, to investigate how CA may modulate mucosal responses to helminth infection [20]. We show that CA regulates mucosal inflammatory responses and enhances immunity to *H. polygyrus*, suggesting that CA-containing dietary supplements may play a role in gut homeostasis and control of intestinal pathogens.

## 2. Materials and Methods

### 2.1 Mice

All experimentation was approved by the Experimental Animal Unit, University of Copenhagen, and conducted according to the guidelines of the Danish Animal Experimentation Inspectorate (Licence number 2015-15-0201-00760). Female C57BL/6 mice (6-8 weeks of age; Janvier, France) were housed at the Department of Experimental Medicine, University of Copenhagen, in individually ventilated cages with ad libitum access to standard mouse chow (D30, SAFE). Mice were administered CA (Sigma-Aldrich; 25 μg/mL) in the drinking water. CA was solubilized in polysorbate-80, and control mice received drinking water with an equivalent amount of polysorbate-80 alone. Mice received the CA-supplemented water for 21 days. Water was refreshed every 2 days and mouse body weights were recorded weekly. For infection experiments, mice were infected with *H. polygyrus* after 7 days of CA treatment and sacrificed by cervical dislocation at day 14 post-infection. Jejunum tissues were transferred to RNAlater (Thermo Fisher) and stored at −20 °C. Digesta samples from the cecum were collected and stored at −80 °C until needed.

### 2.2 Parasites

*H. polygyrus* was propagated and HES produced as described previously [21]. Mice were infected with 200 third-stage larvae by oral gavage. At necropsy, worm burdens were determined using a dissecting microscope.

### 2.3 Flow Cytometry

Mesenteric lymph nodes (MLN) were isolated from mice as described previously [22]. MLN cells were washed and surface stained with anti-TCRβ-FITC (clone H57-597) and anti-CD4-PerCP-Cy5.5 (clone RM4-5). Intracellular staining was carried out with Anti-T-bet-Alexafluor 647 (clone 4B10), anti-GATA3-PE (clone TWAJ) and anti-Foxp3-FITC (clone FJK-16s). All antibodies were sourced from Thermo Fisher. A BD Accuri C6 flow cytometer was used for analysis.

### 2.4 Cell culture

Murine rectal epithelial CMT93 (ATCC-CCL-223) cells or RAW 264.7 macrophages (ATCC-TIB-71) were cultured in DMEM medium with 10% fetal calf serum, 100 U/mL penicillin and 100 μg/mL streptomycin. For stimulation of CMT93 cells, cells of approximately 80% confluence were incubated with CA for 6 hours prior to RNA extraction. RAW264.7 cells (2.5×10 cells/well) were split into 96 well plates for 2 hours to allow cell attachment, followed by simulation with CA for 24 hours together with LPS (500 ng/mL). In some experiments, the Nrf2 inhibitor ML385 (5 μM, Sigma-Aldrich) or *H. polygyrus* Excretory-Secretory antigen (30 μg/mL) were included. After 24 hours stimulation, supernatant or cells were collected for cytokine or ROS analysis, respectively.

### 2.5 RNA extraction and microarray analysis

RNA was isolated from either cell lysate or jejunal tissue using RNeasy kit or miRNeasy mini kits, respectively, according to manufacturer’s protocols (Qiagen, Denmark). Microarray analysis was carried out on tissue and cell RNA as previously described, utilizing the Affymetrix Clariom S HT-24 pipeline [22]. The data was analysed by Transcriptome Analysis Console software (Thermo Fisher), and Gene set enrichment analysis (GSEA; Broad Institute) was used for pathway analysis. Microarray data is available at GEO under accession number GSE165377.

### 2.6 Quantitative Real Time PCR

RNA was extracted as above, and QuantiTect Reverse Transcription Kit (Qiagen) was used for cDNA synthesis. Primers are listed in Table 1. qPCR was conducted with PerfeCTa SYBR Green FastMIX Low ROX (Quanta Bioscience). The qPCR protocol was an initial denaturation step at 95 °C for 2 minutes followed by 40 cycles of 15 seconds at 95 °C and 20 seconds at 60 °C. GAPDH was used as a reference gene for normalization.

**Table 1.**
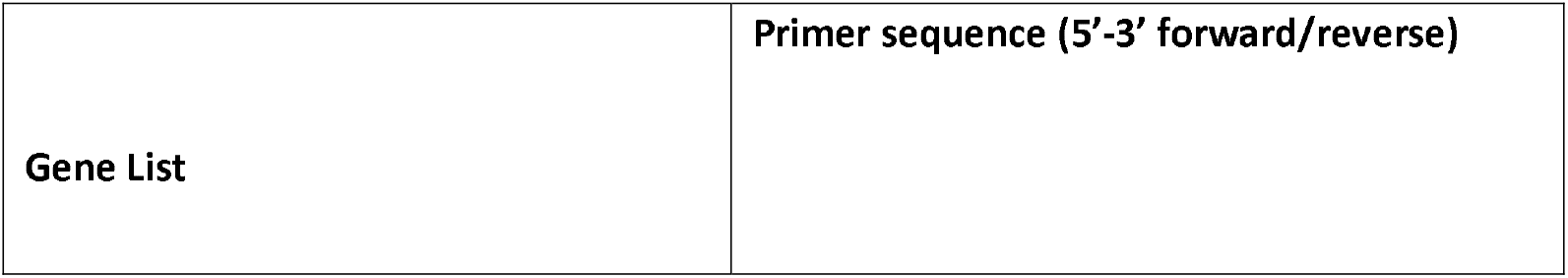
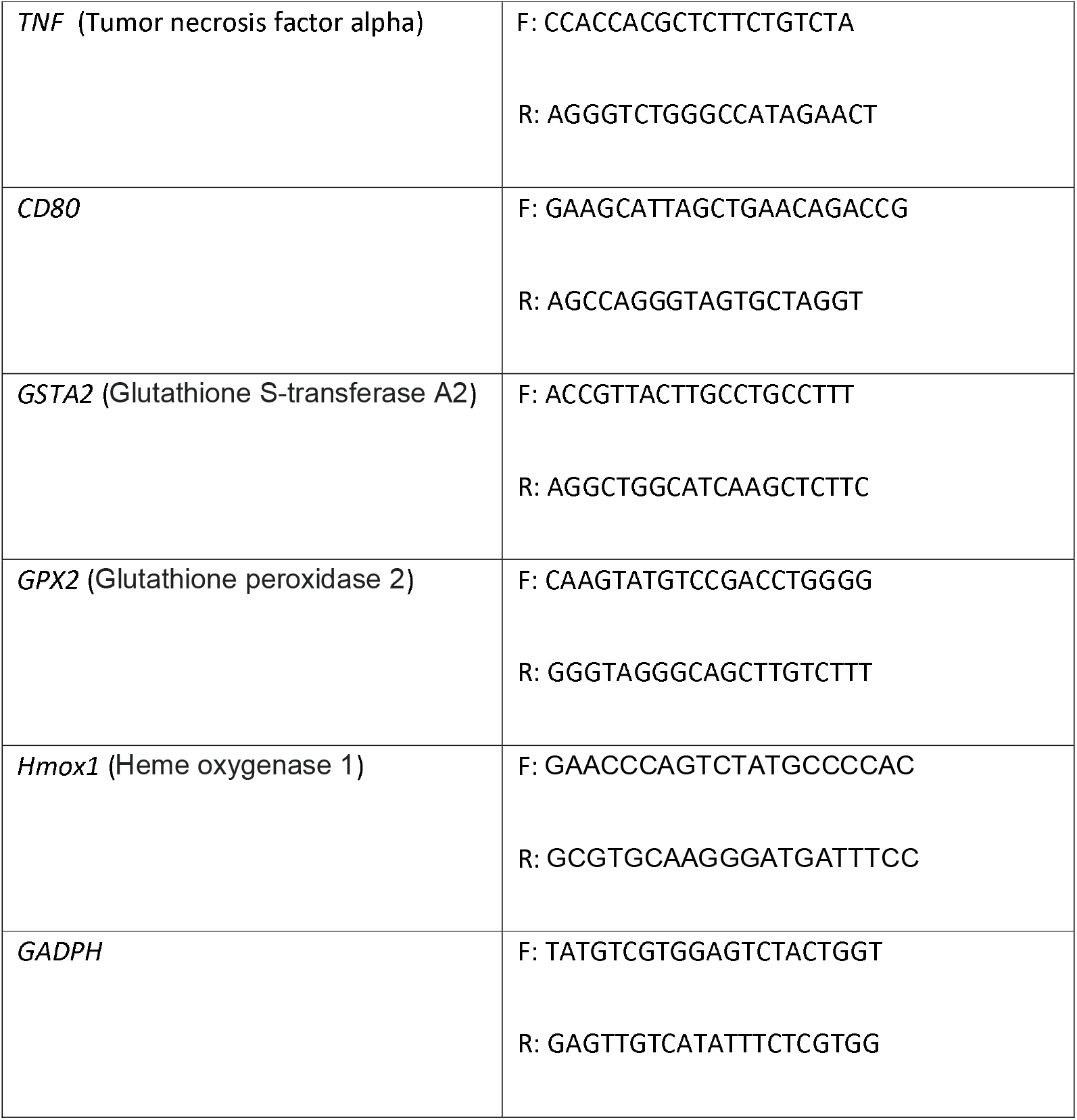

### 2.7 ELISA

IL-6 secretion from RAW264.7 cells was measured using the mouse IL-6 Duoset ELISA kit (R&D systems, UK) according to manufacturer’s procedure.

### 2.8 ROS Measurement

Cells were collected as described above and washed with warm serum-free DMEM. DCFH-DA (10 μM, Sigma) was used to stain cells for 30 min at 37°C. Then DCFH-DA was removed by washing 3 times, before measurement of fluorescence intensity by flow cytometry (Accuri C6, BD biosciences).

### 2.9 Sequencing of cecal microbiota

DNA was extracted from 100mg of cecal contents using the Bead-Beat Micro AX Gravity kit (A&A Biotechnology, Poland) according to the manufacturer’s instructions with addition of mutanolysin and lysozyme to enhance bacterial cell wall degradation. Amplification of the V3 region of the 16S rRNA gene and sequencing was conducted as previously described [22]. The raw dataset containing pair-ended reads was merged and trimmed, and zero-radius operational taxonomic units (zOTU) obtained using the Greengenes (13.8) 16S rRNA gene collection as a reference database. The Quantitative Insight Into Microbial Ecology open source software package QIIME 2 (v2019.4.0) was used for subsequent analysis. Prior to alpha- and beta-diversity analysis samples were normalised to 6000 reads/sample which constituted to the 80% of the most indigent sample in the dataset. Permutational multivariate ANOVA (PERMANOVA) was used to evaluate group differences based on weighted and unweighted UniFrac distance matrices.

### 2.10 Statistical analysis

Statistical analysis of *in vivo* data was conducted using general linear model (SPSS Statistics 27). The general model included fixed factors of diet and infection with blocking by experiment where appropriate. For *in vitro* data T-tests and one-way ANOVA were used (Graph Pad Prism Version 7.05). All data are represented as means ± standard error of mean (SEM). *P* value < 0.05 was considered significant.

## 3. Results

### 3.1 Cinnamaldehyde suppresses transcriptional pathways related to inflammation in the intestine

To gain a global overview of the mucosal response to CA intake, we first assessed the response of small intestinal tissue after 21 days of CA intake in healthy, uninfected mice using a transcriptomic approach. Microarray analysis indicated that mice administered CA clustered distinctly from control mice (Figure 1A). Amongst the genes most significantly down-regulated by CA, were genes related to antigen presentation, such as *h2-dma* and *cd80* (Figure 1B). Down-regulation of *cd80* and *tnf* was verified by qPCR (Figure 1C). Gene pathway analysis showed clearly that the most down-regulated pathways were nearly all connected to inflammatory responses. Notably, TLR and T-cell receptor signalling were amongst the most affected pathways (Figure 1D). Gene pathways that were upregulated by CA were related mainly to biological process connected to olfactory responses and hormonal changes (Figure 1E). Thus, consumption of CA is associated with distinct mucosal changes in the small intestine that mainly relate to immune function and inflammation.

**Figure 1.**
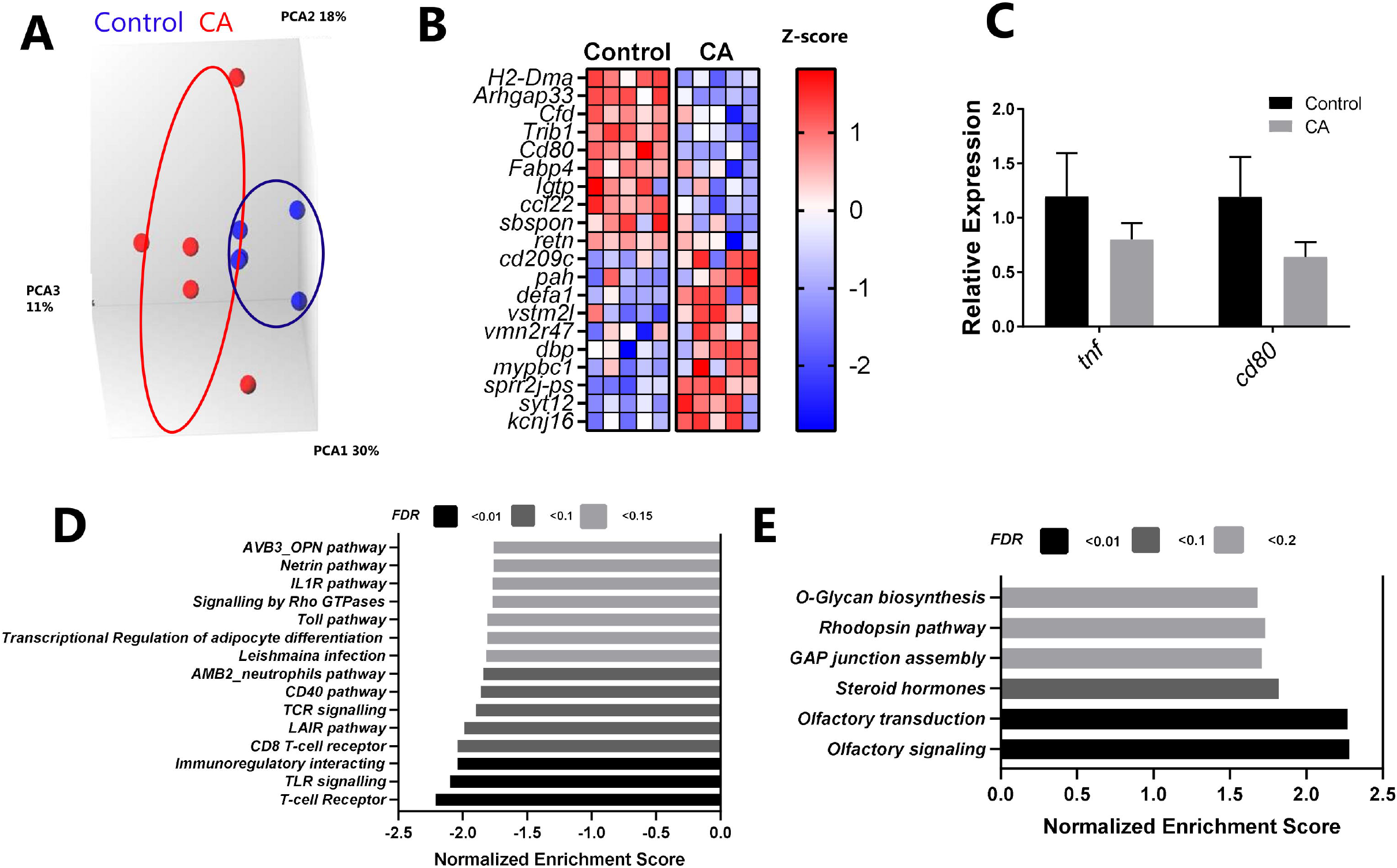
Cinnamaldehyde (CA) modulates transcriptional pathways in the mouse jejunum. **(A)** Principal component analysis (PCA) of gene expression assessed by microarray after 21 days CA uptake. **(B)** Top up- and down-regulated genes by CA treatment **(C)** Relative expression of TNF and CD80 by qPCR. **(D-E)** Significantly down-regulated **(D)** and up-regulated **(E)** gene pathways due to CA intake, identified by gene-set enrichment analysis, grouped by Enrichment Score and FDR (false discovery rate). n=5 mice per group.

### 3.2 Cinnamaldehyde regulates intestinal parasite infection and T-cell populations in mesenteric lymph nodes

Given the significant ability of CA to modulate mucosal immune pathways, we next asked if CA could modulate the response towards an enteric helminth infection in mice. We administered mice CA in drinking water prior to and throughout a 14-day infection period with *H. polygyrus*. The effect of CA on weight and worm burdens are shown in Figure 2. Within infected mice, CA treatment tended to increase growth rates (p=0.075; Figure 2A). Moreover, worm burdens were significantly reduced at 14 dpi (p<0.05; Figure 2B). To explore if CA enhanced the development of Th2 responses, which are protective against *H. polygyrus* infection, we examined mesenteric lymph node cells for proportions of Th1 (T-bet^+^), Th2 (GATA3^+^) and Treg (Foxp3^+^) CD4^+^ T-cells. As shown in Figure 2C, *H. polygyrus* infection significantly increased the proportion of Th2 and Treg cells, with no effect on Th1 cells. CA did not influence Th1 cell numbers, but tended to promote Th2 cells in uninfected mice, although this was less evident in infected mice. Consistent with this, CA significantly enhanced the Th2/Th1 ratio in uninfected mice, and to a lesser extent in infected mice (p<0.05 for interaction between diet and infection), indicating that CA could favour the development of Th2 immunity at the expense of Th1 responses. CA also slightly increased the proportion of Tregs, but this was not significant (*p*=0.2). Thus, CA treatment significantly promotes Th2 type responses in parasite-naive mice, and enhances resistance to *H. polygyrus* infection, albeit without a clear modulation of helminth-induced T-cell responses.

**Figure 2.**
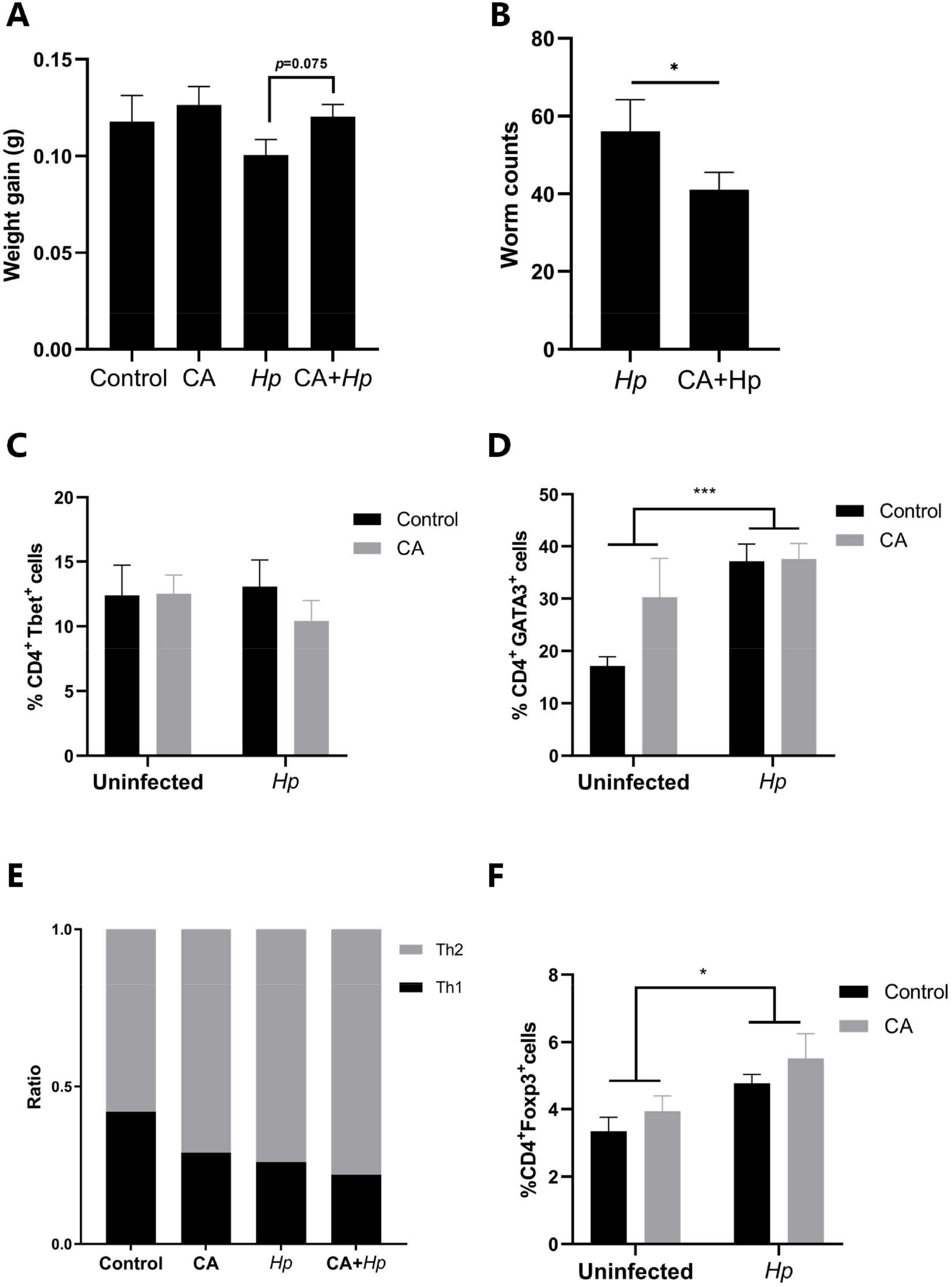
Cinnamaldehyde (CA) modulates the immune response to *Heligomosomoides polygyrus* (*Hp*) infection. **(A)** Average weight gain per day of mice **(B)** Worm burdens in mice fed either CA or control animals, (C-E) Percentage of CD4^+^Tbet^+^ **(C)** CD4^+^GATA3^+^ **(D)** and the ratio of CD4^+^GATA3^+^/CD4^+^Tbet^+^ T-cells **(E)** in mesenteric lymph nodes. Percentage CD4^+^Foxp3^+^ **(F)** T-cells in mesenteric lymph nodes. Data is from three **(A,B)** or two **(C-F)** independent experiments, each with 5 mice per treatment group. Data bars represent mean ± SEM. *p≤0.05, **p≤0.01.

### 3.3 Cinnamaldehyde modulates the mucosal transcriptomic response to *H. polygyrus* infection

To gain further insight into the mechanism underlying the beneficial effects of CA during *H. polygyrus* infection, we conducted transcriptomic analysis on tissues harvested from infected mice with or without CA supplementation. In control-fed mice, *H. polygyrus* infection induced upregulation of characteristic helminth-related genes and gene pathways, relative to uninfected mice. These included genes encoding mast cell proteases (e.g. *mcpt1*), and gene pathways connected to arachidonic acid metabolism, IgE production and asthma, typical of a Th2 polarized immune response (Figure 3A-B). Interestingly, in control-fed mice, infection significantly down-regulated intestinal gene expression pathways related to mitosis and cell cycle, relative to uninfected mice (Figure 3B). When we compared the transcriptomes of *H. polygyrus*-infected mice from the control or CA-fed groups, we found that CA had a profound effect, with the two dietary groups forming distinct clusters, as previously observed in uninfected mice (Figure 3C). Relative to control-fed infected mice, gene pathways enriched in infected mice administered CA were mainly related to cell cycle, mitosis, and DNA replication, indicating that CA restored the mitotic pathways suppressed by infection (Figure 3D). Inspection of genes that were significantly up-regulated by CA in *H. polygyrus*-infected mice showed a number of genes involved in humoral immune and antioxidant responses (*cfd, cr2, Nt5e, Hp*). Conversely, CA suppressed the *H. polygyrus*-induced up-regulation of inflammatory-related genes such as *rasgef1b* and *fos* (Figure 3E). Thus, consistent with its role as an immunomodulatory agent, CA appeared to regulate inflammatory responses to reduce harmful inflammation and restore intestinal homeostasis disrupted by parasite infection.

**Figure 3.**
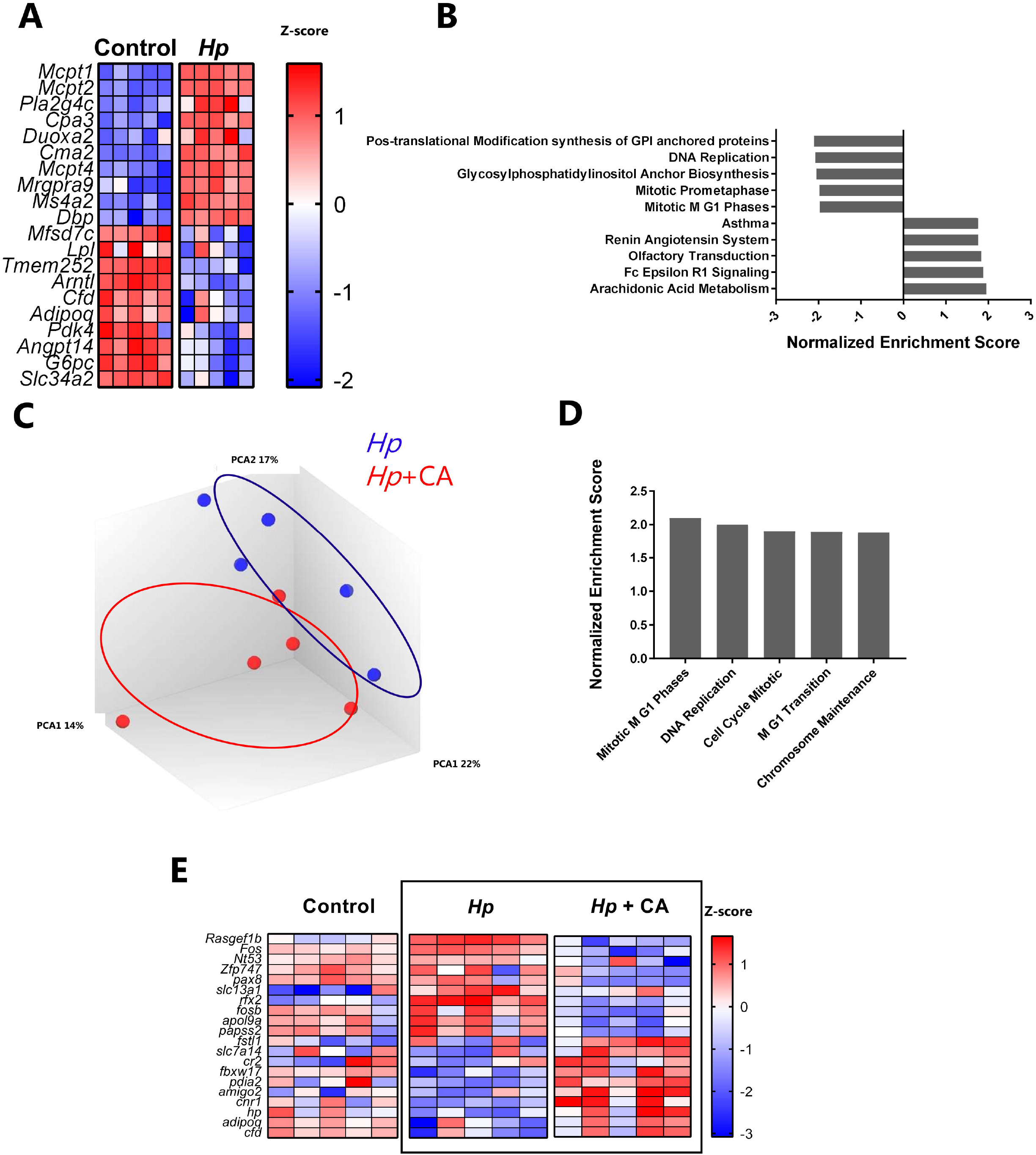
*Heligomosomoides polygyrus* (*Hp*) infection induces changes in jejunal tissue which can be modulated by cinnamaldehyde (CA). **(A)** Top regulated genes resulting from 14-day *Hp* infection in control-fed mice. **(B)** Top regulated pathways resulting from *Hp* infection in control-fed mice, identified by gene-set enrichment analysis, grouped by Normalized Enrichment Score (p<0.001; FDR<0.2). **(C)** Principal component analysis (PCA) of gene expression assessed by microarray after 14 days *Hp* infection in control-fed mice or mice administered CA. **(D)** Top up-regulated gene pathways in *Hp*-infected mice administered CA, relative to control-fed mice infected with *Hp*, identified by gene-set enrichment analysis, grouped by Normalized Enrichment Score (p<0.001; FDR<0.05). **(E)** The most up- and down-regulated genes as a result of CA intake in *Hp*-infected mice. Shown are top ten up- and down-regulated genes identified by comparison of control-fed, *Hp*-infected mice (‘*Hp*’) and CA-fed, *Hp*-infected mice (*Hp* + CA), as well as the corresponding expression in uninfected, control-fed mice (‘Control’). n=5 mice per group.

### 3.4 CA exerts minor changes in the gut microbiota composition

Dietary substances can interact dynamically with the GM, and changes in GM composition can regulate intestinal immune responses to infection [23, 24]. Thus, we next examined whether CA intake altered the composition of the GM, and thus whether a prebiotic effect may be responsible for the immunomodulatory activity. GM composition in cecal digesta samples was determined by 16S rRNA gene amplicon sequencing of the V3 hypervariable region. Analysis of α-diversity demonstrated no effect of CA in uninfected mice (Figure 4A). In the absence of CA, mice infected with *H. polygyrus* had a less diverse GM than uninfected mice (P=0.08), however this effect of infection was not evident in CA-fed mice (P>0.3; Figure 4A). Analysis of β-diversity by unweighted UniFrac showed similar trends. In uninfected mice, CA had no clear effect on GM composition (Figure 4B). Similar to α-diversity, *H. polygyrus* infection resulted in a divergence from uninfected mice (P=0.08 after multiple comparison testing), with this divergence being less evident when CA was administered (P=0.23; Figure 4B). The most pronounced effect of *H. polygyrus* was a marked reduction in the abundance of the phylum Bacteroidetes and an increase in Firmicutes (P<0.001). This could be mainly explained by a reduction in the *Muribaculaceae* (also known as S24-7) family within Bacteroidetes, and an increase in the relative abundance of genus *Allobaculum* within Firmicutes; the magnitude of both these differences was lower in infected mice fed CA (Figure 4C-E). No significant differences at the phylum level were observed as a result of CA intake, although we noted that CA increased the abundance of the Clostridiales order and suppressed the abundance of the genus *Coprobacillus* (Figure 4F-Figure 4G). Clostridiales, an important contributor of amino acid metabolism, has been correlated with IFNγ and regulatory T cells expression, while *Coprobacillus* is associated with serum leptin positively [25–27]. Collectively, these data show that *H. polygyrus* tended to change the cecal microbiota and that CA appeared to attenuate these parasite-induced changes, consistent with the worm burden and transcriptomic data indicating a limited effect on the pathological response to infection. However, the minor effects of CA alone on the GM composition suggest that an independent prebiotic capacity of CA is unlikely to underlie its immunomodulatory effects.

**Figure 4.**
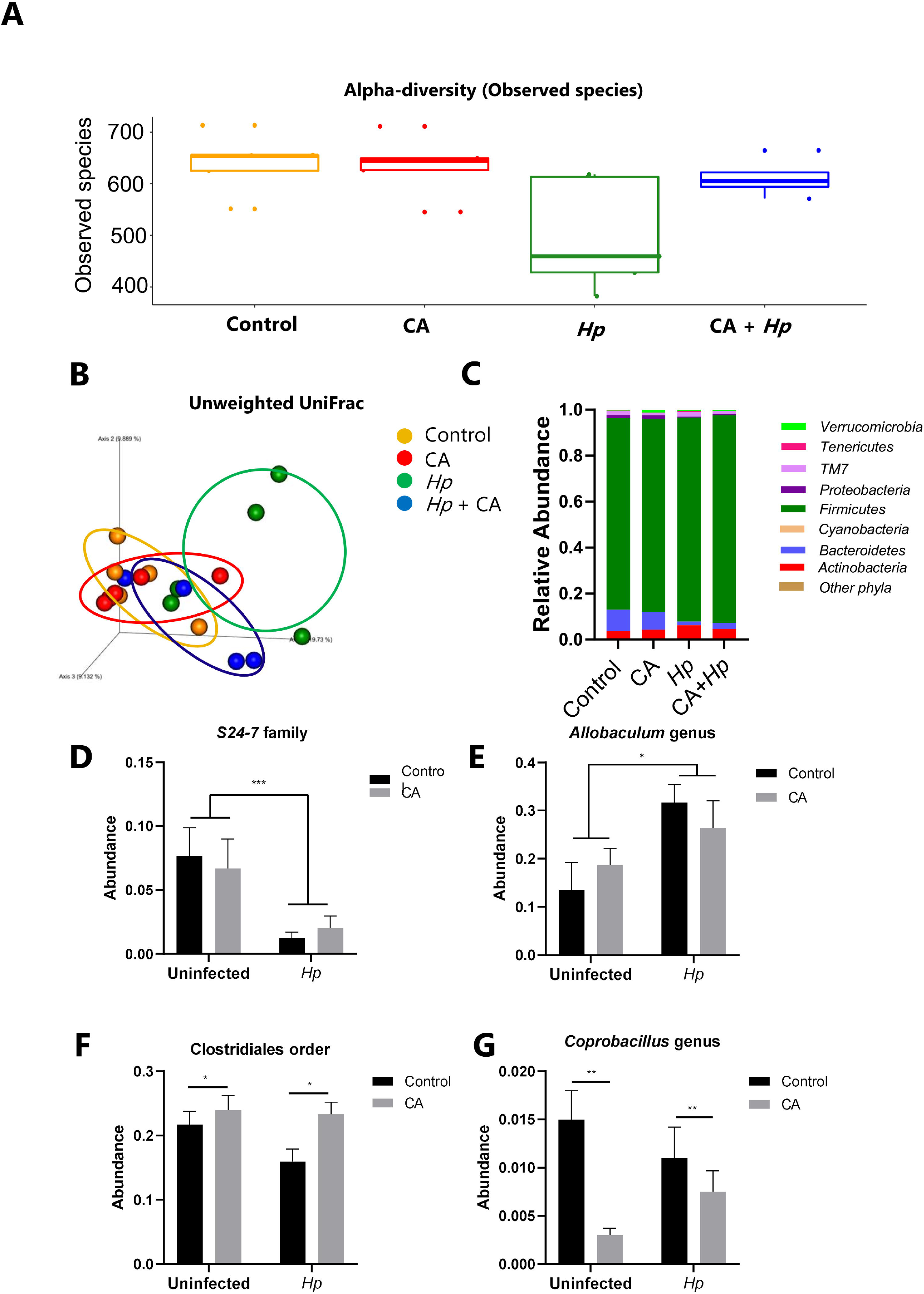
Effects of cinnamaldehyde (CA) and *Heligomosomoides polygyrus* (*Hp*) infection on composition of the cecal microbiota. **(A)** Alpha diversity rarefaction curves showing trend (p=0.08) for reduced diversity in mice infected with *Hp* relative to control mice, but not in *Hp*-infected mice fed CA. **(B)** β-diversity by unweighted UniFrac showing a divergence in mice with different treatment. **(C)** Relative abundance of cecal bacteria at phylum level. **(D)** Abundance of unnamed genus, *S24-7* family. **(E)** Abundance of *Allobaculum* genus. **(F)** Abundance of unnamed genus, *Clostridiales* family. **(G)** Abundance of *Coprobacillus* genus. Data bars represent mean ± SEM. *p≤0.05, **p≤0.01. n=5 mice per group.

### 3.5 CA regulates xenobiotic-metabolizing pathways in intestinal epithelial cells and suppresses inflammatory responses in macrophages *in vitro*

As CA did not substantially modulate the GM, we hypothesized that the immunomodulatory effects derive from direct interactions between CA and host intestinal cells. Therefore, we next investigated the response of intestinal epithelial cells to CA *in vitro*, to gain insight into the mechanistic interactions between CA and the gut tissues. We cultured murine intestinal epithelial cells (CMT93) with CA and assessed transcriptomic responses by microarray. The most upregulated genes were mainly Nrf2-responsive genes involved in xenobiotic detoxification, such as Gsta2 and Gclc as well as the antioxidant genes *Hmox1* and *Aldh3a1* (Figure 5A). Consistent with this, the major pathways induced by CA included glutathione metabolism and signal transduction (Figure 5B). Expression of several of these genes was confirmed by qPCR (Figure 5C). These data suggest that the dominant response to CA exposure in epithelial cells is Nrf2-induced xenobiotic metabolism and antioxidant signalling.

**Figure 5.**
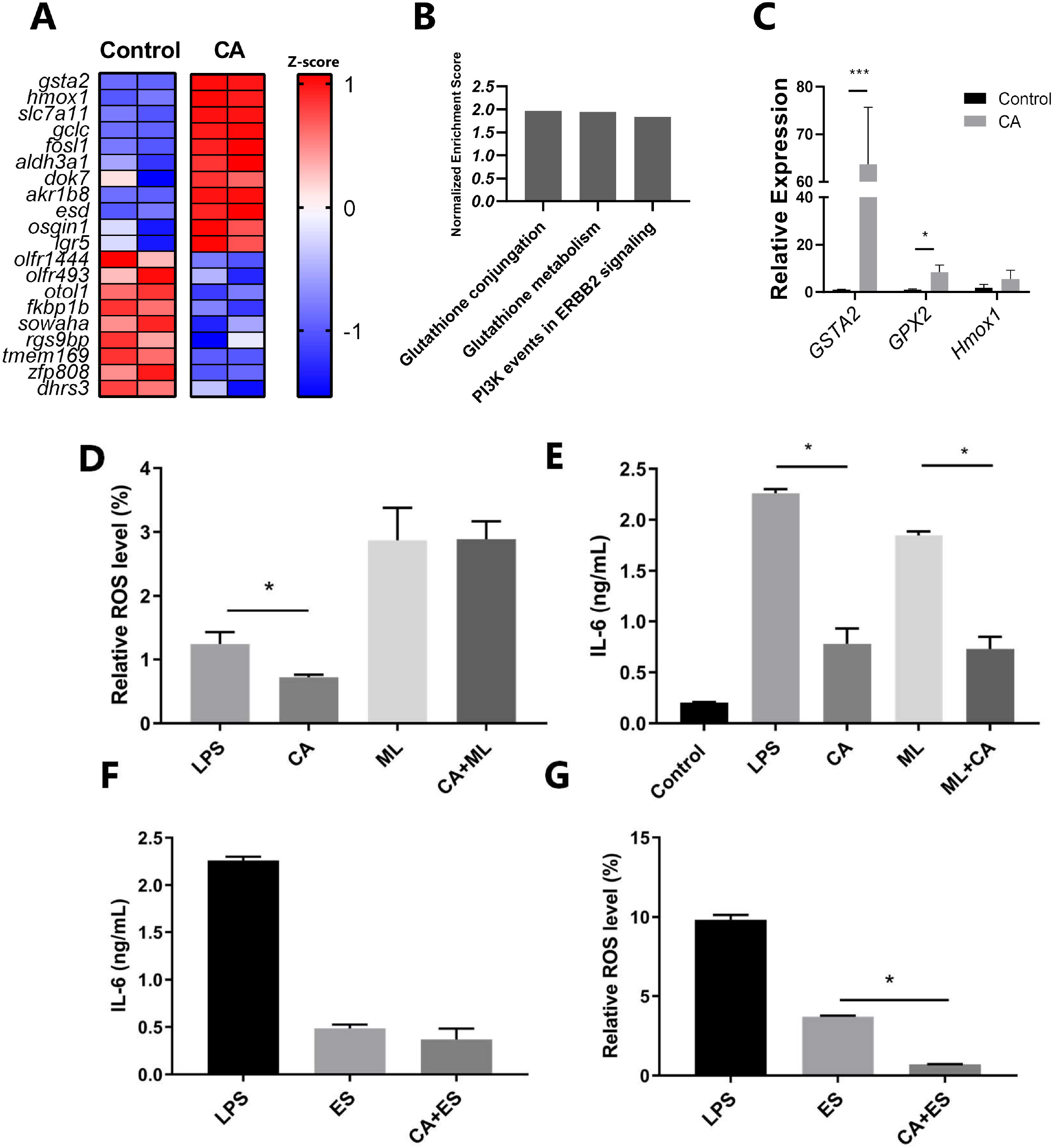
CA stimulation regulates transcriptional pathways and inflammatory responses *in vitro*. **(A)** Top up- and down-regulated genes resulting from stimulation of CMT-93 cells with CA. **(B)** Top up-regulated gene pathways in CMT93 cells stimulated with CA, identified by gene-set enrichment analysis, grouped by Normalized Enrichment Score (p<0.001; FDR<0.05). **(C)** Relative gene expression of gsta2, gpx2 and hmox1 by qPCR. **(D)** Relative reactive oxygen species (ROS) production in CMT93 cells stimulated with LPS or LPS+CA with/without Nrf2 inhibitor ML385. **(E)** Cytokine IL6 production in RAW264.7 cells stimulated with LPS or with LPS+CA with/without Nrf2 inhibitor ML385. **(F)** Cytokine IL6 production in RAW264.7 cells stimulated with CA and *H. polygyrus* excretory/secretory antigens (HES). **(G)** Relative ROS production in RAW264.7 cells stimulated with CA and HES. Data bars represent mean ± SEM. *p≤0.05, **p≤0.01, ***p≤0.001. Data is representative from one experiment with duplicate cells (A-C), or at least two experiments with duplicate cells (D-G).

To confirm the antioxidant effect of CA, LPS induced reactive oxygen species (ROS) production was measured in CMT-93 cells with or without CA exposure. CA inhibited LPS-induced ROS production, however this inhibitory effect was abrogated by concurrent incubation with the Nrf2 inhibitor ML385 (Figure 5D). Thus, CA directly regulates antioxidant capacity in epithelial cells. To assess whether CA also directly regulate inflammatory responses, we assessed pro-inflammatory (IL-6) cytokine production in RAW264.7 macrophages after stimulation with LPS. CA strongly suppressed the secretion of IL-6, however, unlike ROS production, Nrf2-inhibition did not prevent the activity of CA (Figure 5E). *In vivo*, IL-6 is known to impair immunity to *H. polygyrus* by impeding Th2 responsiveness; thus, the strength of the IL-6 activity is inversely related to the *H. polygyrus*-specific immune response [28]. To test whether CA may directly interact with *H. polygyrus* antigens, we exposed LPS-activated RAW264.7 cells to *H. polygyrus* excretory/secretory antigens (HES) with or without CA and quantified IL-6 and ROS production. HES suppressed IL-6 production in RAW264.7 cells, similar to the activity of CA (Figure 5F). Moreover, CA synergized with HES to reduce ROS production (Figure 5G). Collectively, these data indicate that CA activates xenobiotic-metabolizing pathways and induces an antioxidant state in intestinal epithelial cells and directly inhibits inflammatory cytokine release from macrophages. Furthermore, CA augments the immunomodulatory capacity of HES *in vitro*, suggesting that CA may directly modulate the immune response to *H. polygyrus* infection.

## 4. Discussion

In this study, we have investigated the immunomodulatory effects of CA in the murine intestine. The results showed that CA down-regulated inflammation-related pathways in healthy mice and in helminth-infected mice. *In vitro*, CA treatment altered Nrf2 related xenobiotic-metabolizing pathways and immune related genes, and decreased LPS-induced production of ROS and IL-6. Thus, our findings may give more insight into the immune-modulating mechanisms of CA as a feed additive during enteric infection.

CA as a dietary supplement is characterized by its anti-inflammatory and anti-oxidant properties, which can ameliorate some dysfunctions such as diabetes and colitis [10, 29]. Our results revealed a clear anti-inflammatory efficacy in the gut of healthy mice with down-regulated pathways such as the IL1R pathway, TLR signalling and LAIR pathway (local acute inflammatory response). We next analyzed whether CA has the ability to attenuate intestinal disease caused by enteric helminth infection. We observed that CA caused anti-helminth activity in C57BL/6 mice. This result is consistent with other anthelmintic studies *in vitro*, but contrast to our previous study in pigs with *Ascaris suum* [30]. CA is rapidly absorbed in the stomach and proximal small intestine, and mostly transformed to cinnamic acid [31, 32]. This may not provide sufficient contact between CA and *H. polygyrus* to develop direct anti-parasitic effects, but it appears to modulate metabolic pathways and intestinal integrity which may affect helminth resistance [30]. To support the anthelmintic activity, T cells from murine MLNs were divided into different subgroups for potential immune mechanisms of CA. *H. polygyrus* displayed strong ability to activate Th2 and Treg immunity, and CA altered the Th2/Th1 ratio indicating CA regulates the Th2/Th1 balance towards to Th2 polarization. CA treatment can also modulate DNA replication and cell cycle pathways, which were down-regulated by infection alone compared with control mice, showing that CA restored these pathways in infected mice. Furthermore, significant down-regulated genes included Fos, Nt5e and Rasgef1b, known as inflammation-related transcription factor, T cell activation related enzyme and TLR-inducible gene, respectively [33–35]. Conversely, CA in infected mice induced up-regulation of Cfd and *Hp*. *Hp* participates in the modulation of IL17 production and reduction of inflammation severity, as well as Hmox1 induction, showing CA driven regulatory effects in mice may also be connected to xenobiotic metabolism found *in vitro* [36, 37]. More experiments are needed to ascertain the efficacy of CA supplementation on *H. polygyrus* infection.

There is increasing evidence that gut microbial communities are closely related to mucosal immunity, regulated by dietary intake, which may be responsible for distinct immunoregulatory changes. As other studies reported, *H. polygyrus* infection changed the cecal microbiota composition with markedly increased Firmicutes and decreased *Bacteroidetes* in C57BL/6 mice [38–40]. In contrast, CA had no significant difference on microbiota composition at phylum level. While CA treatment reduced *Coprobacillus* genus in mice, which is negatively correlated with cholesterol metabolism, supporting the ability of CA on regulation of lipid metabolism [41]. Whilst CA alone had minimal effects on the microbiota composition, we did note that CA appeared to attenuate the *H. polygyrus*-induced changes in the microbiota, consistent with the lower worm burdens in CA-fed mice.

Considering the minimal effects of CA on the gut microbiota, we hypothesized that CA exerts its immunomodulatory activity through direct interactions with host gut cells. We found that CA treatment significantly up-regulated glutathione related pathways and signal transduction pathway *in vitro*. The glutathione S-transferases family (*Gsta1, Gsta2, Gsta4*) consists of xenobiotic-metabolizing enzymes which have been shown to be related to detoxification process of ROS and thereby protect cells from oxidative stress [42]. We also observed significantly decreased expression of *Dhrs3*, indicating CA may have ability to modulate lipid and retinoid metabolism [43]. The rise of *gpx2* and *Hmox1* with CA treatment may exert intestinal immune regulatory effects through the reduction of inflammatory mediators such as IL-10 and IFN-γ [44, 45]. Notably, *gsta2* gene expression was up-regulated, indicating CA may enhance xenobiotic-metabolizing metabolism *in vitro*. ROS can be induced by LPS as an inflammatory mediator, leading to pro-inflammatory responses with different cytokines. We utilized LPS as a stimulator to produce an inflammatory responses for measuring potential effects of CA or HES on ROS and IL-6 production in macrophages. We found that CA significantly decreased IL-6 and ROS production, similarly to HES, showing the anti-inflammatory function of CA in the presence of helminth secretory products. The ability of CA to accentuate the activity of HES indicates a potential direct effect of CA on *H. polygyrus*-induced immune responses, which may explain the limiting effect of CA on *H. polygyrus* infection *in vivo*.

In conclusion, we observed that CA treatment regulated mucosal immunity in mice. CA appeared to drive Th2 immunity, and restored some signalling pathways disrupted by a small intestinal parasite. CA stimulation also attenuated inflammation related pathways and genes *in vitro* with decreased secretion of ROS and IL-6. Further studies should investigate the practical application of CA as beneficial food components to promote gut health.

## Acknowledgements

We thank Mette Schjelde and Denitsa Stefanova for excellent technical assistance. This work was supported by the Lundbeck Foundation (R252-2017-1731) and Independent Research Fund Denmark (7026-0094B). LZ was supported by the China Scholarship Council (Grant 201806910065). The funding bodies had no involvement in study design, data acquisition or decision to publish.

